# Enrichment of hard sweeps on the X chromosome in *Drosophila melanogaster*

**DOI:** 10.1101/2022.05.26.493648

**Authors:** Mariana Harris, Nandita Garud

**Affiliations:** Department of Computational Medicine, University of California Los Angeles, Los Angeles California, USA; Ecology and Evolutionary Biology, University of California Los Angeles, Los Angeles California, USA; Department of Human Genetics, University of California, Los Angeles, California, USA

## Abstract

The characteristic properties of the X chromosome, such as male hemizygosity and its unique inheritance pattern, expose it to natural selection in a way that can be different from the autosomes. Here, we investigate the differences in the tempo and mode of adaptation on the X chromosome and autosomes in a population of *Drosophila melanogaster*. Specifically, we test the hypothesis that due to hemizygosity and a lower effective population size on the X, the relative proportion of hard sweeps, which are expected when adaptation is gradual, compared to soft sweeps, which are expected when adaptation is rapid, is greater on the X than on the autosomes. We quantify the incidence of hard versus soft sweeps in North American *D. melanogaster* population genomic data with haplotype homozygosity statistics and find an enrichment of the proportion of hard versus soft sweeps on the X chromosome compared to the autosomes, confirming predictions we make from simulations. Understanding these differences may enable a deeper understanding of how important phenotypes arise as well as the impact of fundamental evolutionary parameters on adaptation, such as dominance, sex-specific selection, and sex-biased demography.

## Introduction

Adaptation on the X chromosome has attracted significant interest from evolutionary biologists because its dynamics seem to be distinct from that of autosomes. The X chromosome is hemizygous in males, increasing the visibility of new mutations to natural selection on the X and thus potentially subject to different evolutionary dynamics than autosomes. The X harbors many essential genes, including genes responsible for speciation [1,2], fertility [3], sexual dimorphism [4,5], and brain function [6], as well as several genes that are preferentially expressed in one sex [7–10]. In the classic model organism *Drosophila melanogaster*, there is evidence of a faster rate of adaptive evolution on the X [11–13] and sexually antagonistic selection acting on the sex chromosomes [5,14,15], revealing crucial differences in adaptation between the X and autosomes. Thus, by studying differences in the tempo and mode of adaptation between the X and autosomes we may increase our understanding of evolution at a molecular level, particularly in the context of sexual dimorphism, sex-biased demography, speciation, and sex chromosome evolution.

The tempo and mode of adaptation in natural populations more broadly has been long debated. Adaptation can be characterized as gradual or rapid [16–19], and its pace depends on the availability of adaptive mutations. When these mutations are absent or rare before the onset of selection, either because the effective population size (*N_e_*), adaptive mutation rate (*μ_A_*), or their product (*θ_A_* <<1, where *θ_A_* = 4*N_e_μ_A_*) is small, adaptation is expected to be gradual [16,20]. In such a scenario, hard selective sweeps are expected to be common, in which a single adaptive mutation rises in frequency, leaving behind characteristic deep dips in diversity in the vicinity of the adaptive locus [18,19]. By contrast, when there is a large input of mutations due to large census population sizes and/or mutation rates (e.g. *θ_A_* >1) [20], or, standing genetic variation (SGV) is abundant [16], adaptation is expected to be rapid. In such a scenario, soft selective sweeps are expected to be more common, in which multiple adaptive mutations on distinct haplotypes sweep through the population simultaneously, and do not necessarily result in dips in diversity [18,19].

In addition to soft sweeps being generated by a high input of *de novo* mutations or SGV, a third process can increase the prevalence of soft sweeps: dominance shifts, whereby a recessive deleterious mutation becomes dominant and beneficial in a new environment [21]. In this scenario, evolutionary and physiological theories of dominance predict that loss of function mutations are generally recessive while gain of function mutations are generally dominant [22–25]. Muralidhar and Veller [21] argue that one example of such scenario is at the *Ace* locus in *Drosophila,* which encodes for the enzyme acetylcholinesterase that catalyzes the breakdown of the neurotransmitter acetylcholine and has evolved adaptations in response to pesticides [26–30]. Without pesticides, mutations at the *Ace* locus are deleterious and result in less efficient binding of acetylcholine [29,30]. With pesticides, however, mutations at *Ace* are beneficial because they confer resistance to pesticides [27,31]. Previous work in multiple species has shown that the beneficial effect of pesticide resistant alleles is dominant [32,33], and that the deleterious effect of such mutations in the absence of pesticides is at least partially recessive [31,34,35].

When mutations are slightly deleterious and recessive, their effect on fitness will be initially masked, making it more likely that these mutations can segregate at some low frequency in the population [36–38]. This in turn will increase the number of copies of the variant present in the population when the environmental change occurs, resulting in more distinct haplotypes present in the population at the onset of positive selection. Additionally, with dominance shifts, adaptive mutations in the new environment are expected to be at least partially dominant, and thus are less likely to be lost than if they were still recessive. By this logic, soft sweeps are more likely than hard sweeps when there are dominance shifts as compared to when dominance remains constant across environments [21].

While soft sweeps have been found to be common on the autosomes [39–41], should they be equally common on the X? The differences in the inheritance patterns of the X chromosome and the autosomes, as well as the exposure of mutations on the hemizygous X can give rise to differences in the signatures of selection found on the X compared to those from the autosomes. To begin with, the effective population size of the X, *N_eX,_* is usually expected to be lower than that of the autosomes, *N_eAuto_.* Particularly, in a population with an equal number of males and females there are only three X chromosomes per every four autosomes, hence *N_eX_* _=_ ¾ *N_eAuto_.* This lower population size can increase the effect of genetic drift and also, lower the mutational input on the X such that *θ_A_X__* = 0.75*θ_A_auto__* [42,43]. Moreover, because mutations on the X in males are immediately exposed to selection, recessive beneficial mutations on the X are less prone to be lost due to stochastic forces and recessive deleterious mutations are more likely to be efficiently purged from the population compared to recessive mutations on the autosomes [11,43,44]. All these factors can increase the likelihood of hard sweeps on the X chromosome, either through a lower mutational input from the reduced *N_eX_* or due to a more efficient purifying selection that decreases the genetic variation that can seed adaptation on the X.

In this paper we examine the relative proportion of hard versus soft sweeps on the X and autosomes using the model organism *D. melanogaster*. To date, while evidence for more rapid evolution on the X has been documented in *D. melanogaster*, the relative proportions of hard versus soft sweeps on the X versus autosomes have not been evaluated with a systematic scan. We focus on *D. melanogaster* because the molecular basis of evolution has already been extensively studied in this organism and there exist several well-documented cases of adaptation across the literature. On the autosomes, three cases of recent adaptation are at the loci *Ace*, *Cyp6g1*, and *CHKov1*, due to resistance to pesticides, DDT, and viruses [26–28,45–48]. These three cases were discovered by empirical means and are all soft sweeps arising from either *de novo* mutations or standing genetic variation. On the X chromosome, the gene *Fezzik* is known to be under positive selection as well [14,49], and may experience sexual antagonism. This too was discovered by empirical means, but it is unknown if there is a hard or soft sweep at this locus. To quantify hard and soft sweeps, we used haplotype homozygosity statistics we recently developed [39,50] that are capable of detecting and differentiating both types of sweeps and can recover known positive controls. In previous work, we showed that application of these statistics to the autosomal data in the Drosophila Genetic Reference Panel (DGRP) data set [51], which consists of 205 fully phased genomes from a population in North Carolina, provide evidence for abundant soft sweeps on the autosomes [39,52]. Now, our simulations and application of these same statistics to the X chromosome provide evidence that the proportion of hard sweeps relative to soft sweeps is enriched on the X chromosome.

## Results

We first examined the expected prevalence of hard and soft sweeps on the X versus autosomes in simulations with parameters relevant to *D. melanogaster*. To do so, we used the forward in time simulator SLiM 3 [53,54], which supports simulations of both autosomal and X chromosome evolution (Methods). Next, we examined the incidence of hard and soft sweeps in DGRP data. To do so, we applied haplotype homozygosity statistics we previously developed for detection of hard and soft selective sweeps.

### 1. Simulations of hard and soft sweeps on the X versus autosomes

#### 1.1 Expected prevalence of hard vs soft sweeps as a function of *θ_A_* and dominance coefficient

To understand the expected incidence of hard and soft sweeps on the X versus autosomes, we performed simulations of selection under a wide range of evolutionary scenarios. We varied *θ_A_* given its role in generating hard versus soft sweeps [19,20,55], where *θ_A_X__* = 0.75*θ_A_auto__*. We also varied dominance (*h*) given differences in hemizygosity on the X versus autosome, as well as its recently discovered role in generating hard versus soft sweeps [21]. We defined the softness of a sweep by the number of distinct mutational origins at the locus under selection at the time of fixation in a sample of *n* = 100 haplotypes, resembling the sample size of the DGRP data (**Methods**). A simulation was classified as a soft sweep if the sampled population had more than one mutational origin and as a hard sweep if it had a single origin. Finally, because forward in time simulators are computationally intensive when simulating large populations such as *D. melanogaster* (*N_e_*~1e6), we rescaled the simulation parameters, as described in the Methods.

In agreement with theoretical expectations [16,20], **Figure 1** shows that the number of origins of a sweep increases with *θ_A_* on both the X and the autosomes. While sweeps on autosomes typically have a higher number of origins compared to the X, this difference depends on the dominance coefficient of mutations. When *h*=0, selective sweeps are softer on the X than on the autosomes, whereas for *h*>0, sweeps on the autosomes are softer. As *h* increases, the softness of sweeps on both autosomes and the X chromosome increases. Moreover, the average number of generations that it takes for a sweep to reach fixation on the X is lower than that observed on the autosomes when *h<*0.5 but higher when *h*>0.5 (**Figure S1a**). This is consistent with the Faster-X theory [13,56,57] and with the fact that when adaptation is gradual, selective sweeps signatures are expected to be hard, but if adaptation is rapid signatures of soft sweeps are more likely to arise.

**Figure 1.**
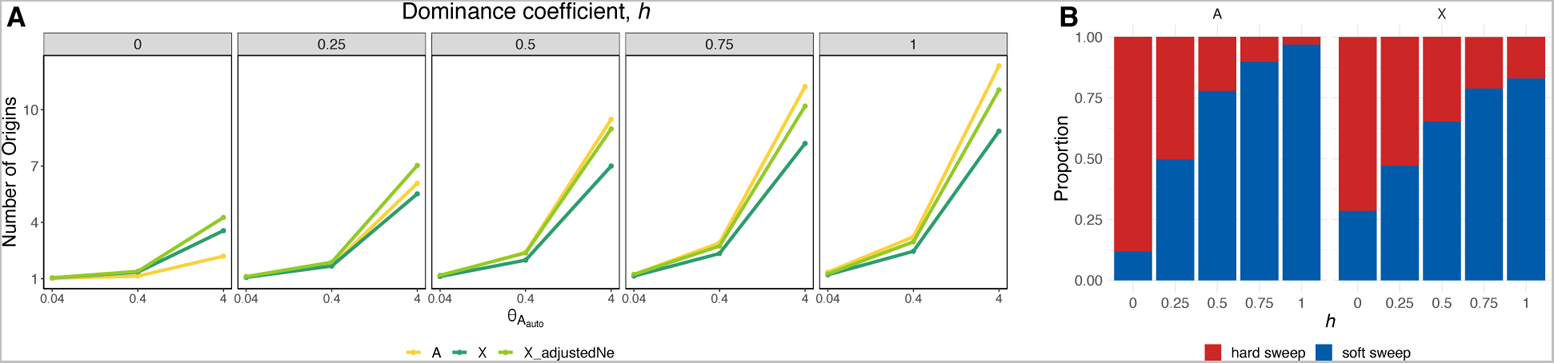
Number of origins as a function of θ_A_ and dominance coefficient. (A) Average number of distinct mutational origins in simulations of selective sweeps from recurrent mutations for *θ_A_* = 0.04, 0.4 and 4 with *N_e_s* =100. The yellow line represents the number of origins for sweeps on the autosome, while the dark green represents the number of origins for sweeps on the X chromosome, in which *θ_A_X__* = 0.75*θ_A_auto__*. The light green line corresponds to simulations in which *N_eX_* was adjusted such that *θ_A_X__* = *θ_A_auto__*. Each panel corresponds to a different dominance coefficient, with *h*>0.5 indicating dominant or partially dominant mutations and *h*<0.5 recessive or partially recessive mutations. (B) Proportion of simulations resulting in hard (red) or soft (blue) sweeps for *θ_A_auto__* = 0.4. For each combination of parameters we ran 1,000 simulations.

To assess the relative contribution of hemizygosity versus lower *θ_A_* in generating harder sweeps on the X, we adjusted *N_eX_* such that *θ_A_X__* = *θ_A_auto__* (light green line in **Figure 1**). In this scenario, the average number of origins increases compared to the un-corrected *N_eX_* (dark green), since increasing *N_eX_*, increases the mutational input and consequently, the probability of soft sweeps. Nonetheless, sweeps on the autosomes appear softer than on the X even with the adjusted *N_eX_* for *h* > 0, indicating that hemizygosity contributes to harder sweeps on the X. In sum, selective sweeps are more likely to be hard on the X chromosome than on the autosomes due to a combination of lower *θ_A_* and hemizygosity.

#### 1.2. Expected prevalence of hard versus soft sweeps as a function of dominance shifts

As previously shown [21], dominance shifts in changing environments can result in softer sweeps on the autosomes, and are hypothesized to result in harder sweeps on the X. To further investigate the effect of dominance on the prevalence of hard and soft selective sweeps, we assessed adaptation in response to changes in the environment following the simulation strategy in Muralidhar and Veller [21].

In this scenario, mutations at the locus of interest were initially deleterious with selection coefficient *s_d_* and dominance given by *h_d_*. Mutations were introduced at a rate *μ_del_* according to parameter *θ_del_* = 4*N_e_μ_del_*. After 10*N_e_* generations, the deleterious mutations segregating in the population, if any, became beneficial with selection coefficient *s_b_* and dominance *h_b_*. For each set of parameter values (*s_d_, h_d_, s_b_, h_b_, θ_del_),* we ran a total of 2,000 simulations and recorded the proportion of simulations that resulted in no standing variation, lost sweep, hard sweep, and soft sweep. No standing variation refers to the case in which there are no copies of the deleterious variant in the population at the time of the environmental change and lost sweep refers to the case in which mutations are lost after they become beneficial. We simulated a model with constant dominance across environments (*h_d_ = h_b_)* as well as a model of dominance shifts where we set *h_b_ =*1*-h_d_*.

**Figure 2** shows that selective sweeps are softer on autosomes than on the X, irrespective of whether or not there is a dominance shift. With dominance shifts, this difference becomes more pronounced the stronger the change in dominance. Additionally, with dominance shifts, soft sweeps become more likely on the autosomes, in agreement with previous results [21]. However, on the X chromosome, the effect of dominance shifts is not as strong as in the autosomes resulting in a slight increase in the likelihood of soft sweeps and a larger proportion of hard sweeps. Finally, the proportion of sweeps that are lost before the environmental change is larger on the X than on the autosomes for all values of *h* (light gray region in **Figures 2b,d**). This higher loss of deleterious mutations on the X can be explained by either more efficient background selection or stronger genetic drift due a lower *N_eX_* [42,44,58]. Together, these observations indicate that selective sweeps are more likely to be hard on the X chromosome than on the autosomes in changing environments, with or without dominance shifts.

**Figure 2.**
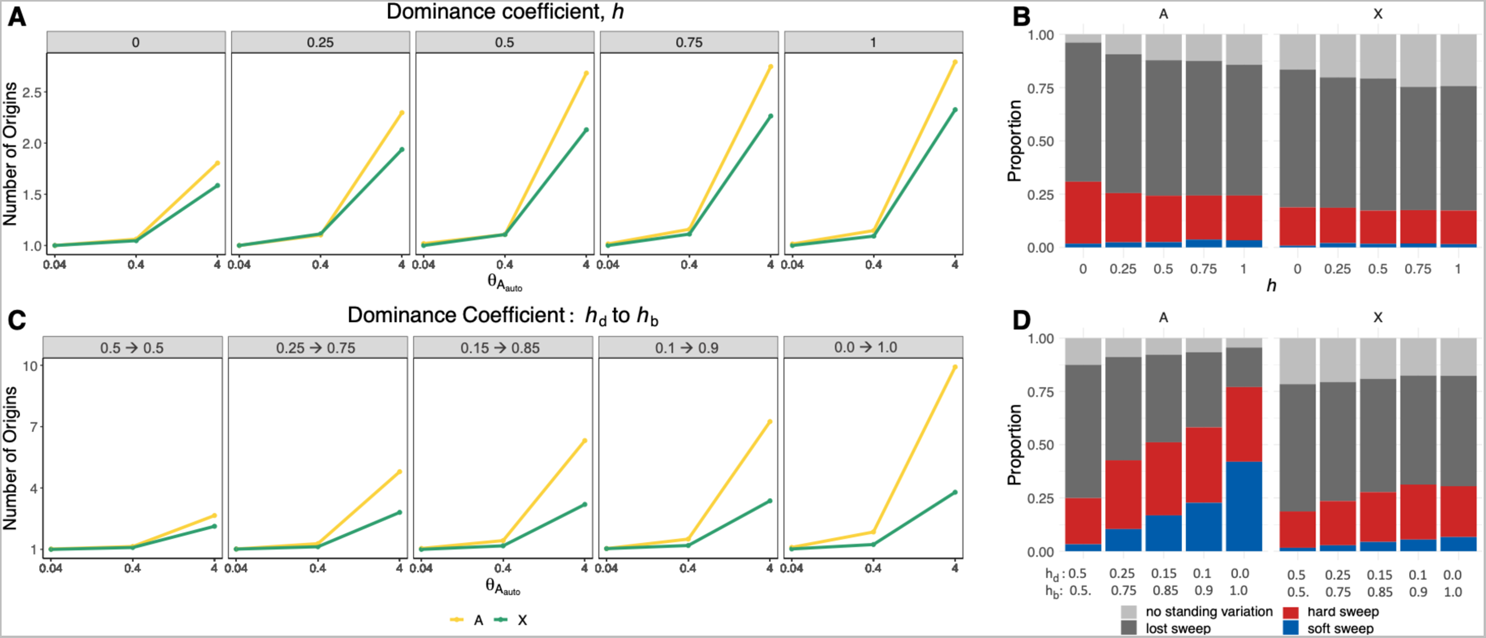
Dominance shifts increase the proportion of soft sweeps on autosomes to a greater extent than on the X. The number of origins on the autosomes and the X for θ_A_ = 0.040, 0.4 and 4 and different dominance coefficients when dominance is constant (*h*) **(A)** or changing (*h_d_* to *h_b_* before and after the onset of selection, respectively) **(C)**. **(B)** The proportion of hard and soft selective sweeps on the autosomes and the X for different dominance coefficients in a model of adaptation to a change in the environment with constant dominance. **(D)** The proportion of hard and soft selective sweeps in the case of dominance shifts. For each combination of parameters, we ran a total of 2,000 simulations of a constant *N_e_*_Auto_=1e6 model, with *N_e_s_d_* and *N_e_s_b_* =100, θ_del_auto__ = 0.4 and θ_del_X__ = 0.75θ_del_auto__.

#### 1.3. Expected prevalence of hard versus soft sweeps as a function of sexual antagonism

Finally, we investigated the effect of sexual antagonism, whereby a mutation can be beneficial for one sex but harmful for the other, as this has been shown to be a common evolutionary force in *D. melanogaster* [14,15,59,60]. The unique inheritance patterns of the X chromosome can lead to sex-dependent selection to act differently between the autosomes and the X. To begin with, autosomes spend equal amounts of time in both sexes, balancing out sexually antagonistic forces, whereas the X chromosome spends 2/3 of its evolutionary time in females and 1/3 in males. This could potentially bias selection on the X to be more favorable for females. However, because of male hemizygosity on the X, selection could also favor mutations that benefit males [4,43,56,61]. Evidence of sexual antagonism influencing genetic variation has been documented in a range of species including humans [10], aphids [7], mice [8], and *Drosophila* [15,49,59,60,62,63]. However, to our knowledge, the influence of sexually antagonistic selection on the prevalence of hard and soft selective sweeps is not well known. Through simulations, we explored how these forces can influence the signatures of selection on the autosomes and the X chromosome.

We simulated two scenarios of sexual antagonism: female-disadvantage with male-advantage and male-disadvantage with female-advantage. To do this, we introduced sexually antagonistic mutations to the population according to parameter *θ_A_* with a beneficial selection coefficient *s_b_* in one sex and a deleterious selection coefficient *s_d_* in the other sex. We set *s_d_* = −*ks_b_*, where *k* is a scalar that modulates the deleterious strength of selection and was set to 0.1. In **Figure 3** we show the number of origins as a function of *θ_A_* and dominance in a scenario where the introduced mutations are deleterious in females but beneficial in males (**Figures 3a-b**) and a scenario where mutations are deleterious in males but beneficial in females (**Figures 3c-d**). In the case of female disadvantage, there is a higher average number of origins on the X when mutations are highly recessive (~*h* <0.25), otherwise the number of origins is lower on the X than on the autosomes. In the case of male disadvantage, there are a lower number of origins on the X for all values of dominance. These observations suggest that under a model of sexual antagonism, selective sweeps are more likely to be harder on the X chromosome than on the autosomes with exception of recessive mutations that are female-deleterious.

**Figure 3.**
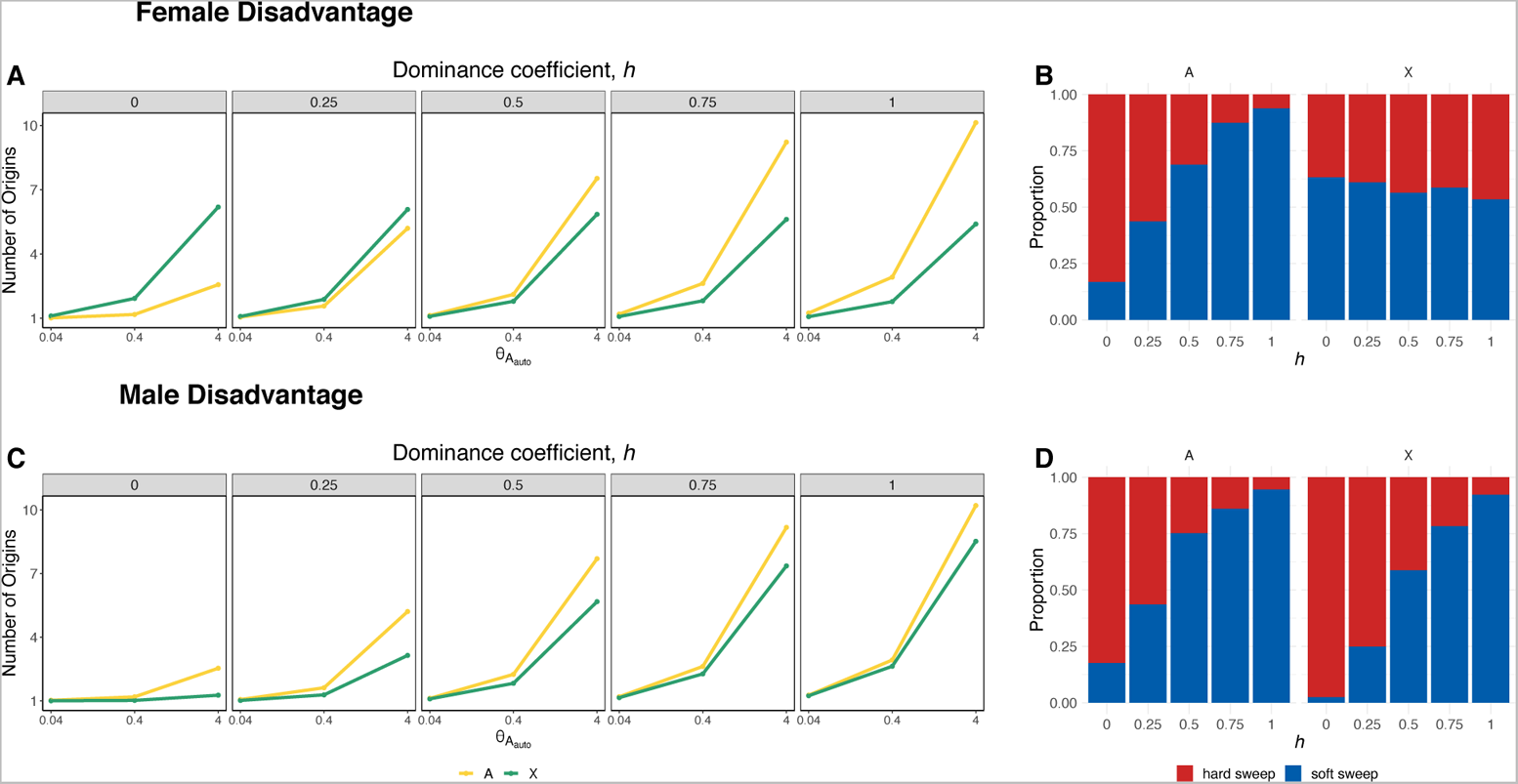
Prevalence of hard and soft selective sweeps for sexually antagonistic selection as a function of θ_A_ and dominance. In (**A-B**) mutations are harmful for females and beneficial in males. Selective sweeps are harder on the X except for the case of strongly recessive mutations (~ *h*<0.25). In (**C-D**) the mutations are deleterious in males but beneficial for females. In this scenario the simulated sweeps are harder on the X for all values of dominance. We ran 1,000 simulations for constant *N_e_*_Auto_=10^6^ model with θ_del_auto__ = 0.04,0.4 and 4 and θ_del_X__ = 0.75θ_delAuto_, *s*_b_=0.01 and *s*_d_=-0.001.

### 2. Analysis of DGRP data

The evolutionary scenarios explored in simulations in the previous section demonstrate that the expected number of origins for selective sweeps on the X chromosome is generally lower than that of autosomes. Next, we examined the prevalence of hard and soft sweeps on the X and autosomes in the DGRP data set [51], composed of 205 inbred *D. melanogaster* genomes from North America.

#### 2.1 Diversity on the X versus autosomes of D. melanogaster

First, we reassessed estimates of S/bp and Pi/bp on the X versus autosomes in two populations of *D. melanogaster*: a derived North American population (DGRP [51]) and an ancestral Zambian population (DPGP3 [64]). A previous study argued that the diversity patterns observed in ancestral and derived genomic data could not be explained by neutral demography alone and proposed a model with a 7:1 female biased ancestral sex ratio combined with a population bottleneck that retained this bias along with higher rates of positive selection on the X chromosome in the derived population [65].

In consonance with the previous findings [66–69], genome-wide diversity is significantly reduced in North America relative to Zambia (p-val < 2.2e-16 one-sided Wilcoxon rank sum test for both for S/bp and Pi/pb; **Figure 4a**), with a more extreme reduction in diversity on the X compared to the autosomes. Based on the average S/bp, we obtain *θ_Zambia_Auto__*/*θ_America_Auto__* = 0.54 (95% confidence interval 0.44-0.64) for autosomal loci and *θ_Zambia_X__*/*θ_America_X__* = 0.31 (0.21-0.41) for the X chromosome. Moreover, S/bp and Pi/bp are significantly reduced on the X chromosome relative to the autosomes in the North American population (p-val < 2.2e-16, one-sided Wilcoxon rank sum test for both for S/bp and Pi/pb; **Figure 4a**), whereas in the ancestral Zambian population, there is no evidence to support a decrease on X chromosome diversity (one-sided Wilcoxon rank sum test p-val=1 and p-val=0.95 for S/bp and Pi/pb, respectively; **Figure 4a**).

**Figure 4.**
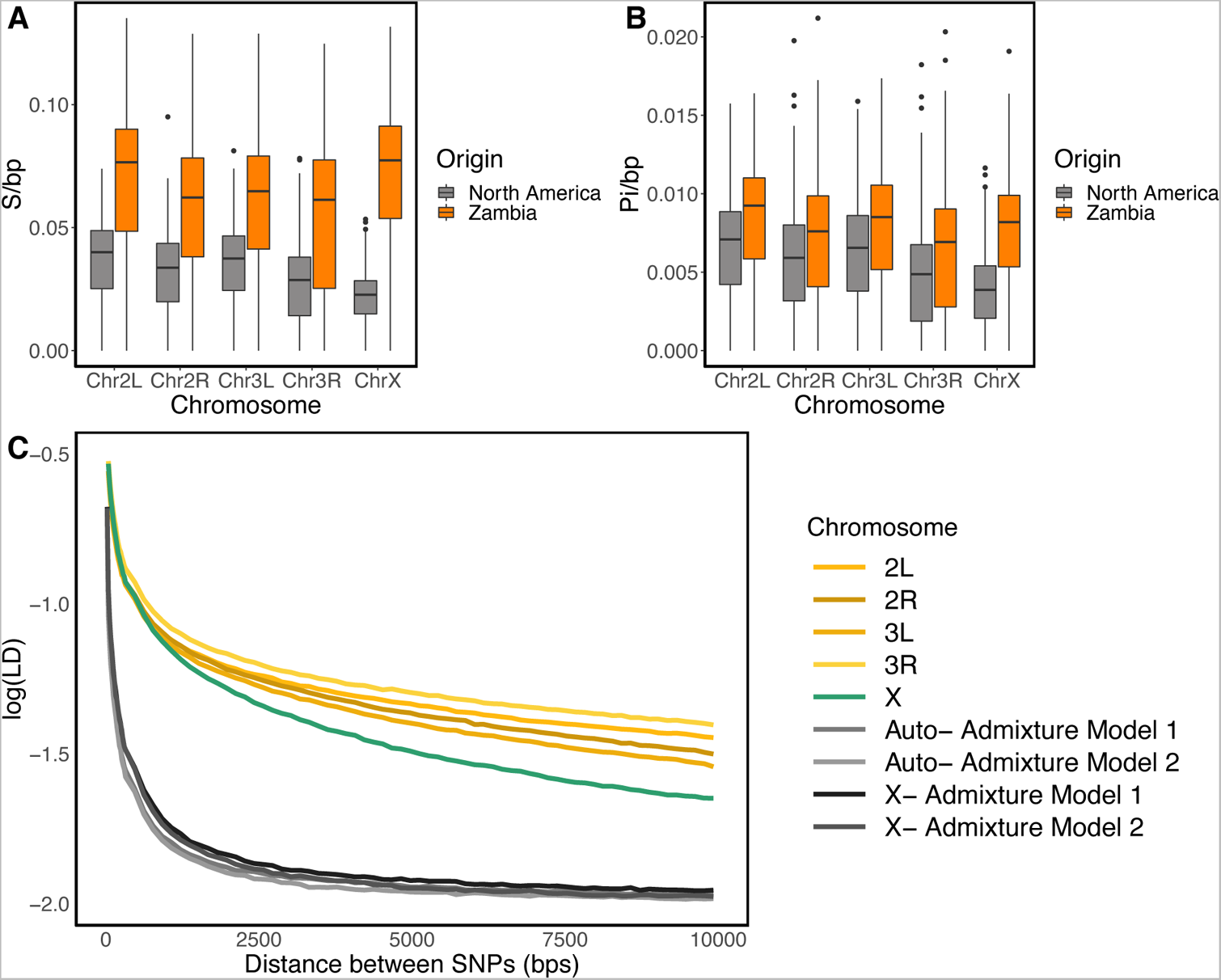
Genetic diversity on the X and autosomes in *D. melanogaster*. S/bp (**A**) and Pi/bp (**B**) in the North American (gray) and Zambian (orange) populations with sample size *n* = 100. (**C**) Pairwise LD in North American data and in neutral admixture Models 1 and 2. Regions with low recombination (ρ≤ 5 × 10^−17^ cM/bp) were excluded from LD computations. Demographic models were simulated with ρ= 5 x 10^−17^ cM/bp (Methods).

To further explore the role of neutral versus selective forces in generating the observed diversity patterns, we next compared levels of linkage disequilibrium (LD) in the North American population with that of two neutral admixture models previously fit to the data [52,70]. These demographic models are variations of the Duchen et al. 2013 admixture model and have been shown to fit the DGRP data in terms of multiple summary statistics [52]. We refer to these models as Model 1 and Model 2, where in Model 1 the contemporary North American population size is *N_eAm_ =1,110,000* and in Model 2 *N_eAm_ = 15,984,500.* The remaining parameters remain the same across the models and are specified in **Table S1** and in [52].

LD in the DGRP data is elevated compared to neutral expectations generated by Models 1 and 2 for both the autosomes and X chromosome (**Figure 4b**). The elevation of LD on the X is notable given overall higher average recombination rates on the X compared to the autosomes (**Figure S2,** [68,71]). We previously argued that the elevated LD on the autosomes is likely due to positive selection [39,52]. Consistent with previous conclusions based on depressed nucleotide diversity on the X, positive selection may also be responsible for the elevated LD on the X. In the next section, we explore the role of positive selection on the X.

#### 2.2 Detection of hard and soft sweeps on the X versus autosomes in DGRP data

To assess the role of positive selection on the X versus autosomes, we next applied the haplotype homozygosity statistic H12, which has the ability to detect hard and soft sweeps [39,52]. To apply H12 to genomic data, one must first define a window size in terms of number of single nucleotide polymorphisms (SNPs). In Garud et al. 2015, 401 SNP windows were used on the autosomal DGRP data, where the average length of these windows (~10kb) was shown to be large enough to avoid detecting regions of high homozygosity due to random fluctuations in diversity, yet not so large that sweeps cannot be detected. With this window size, sweeps with S > 0.05% can be detected.

An H12 scan with 401 SNP windows on the X chromosome shows substantially reduced signal compared to a scan with the same window size on the autosomes (p-val < 2.2e-16, one-sided Wilcoxon rank sum test; **Figures S3 and S4**). Given the elevated LD observed in the data compared to neutral expectations (**Figure 4**) and previous evidence of positive selection acting on the X of *D. melanogaster* [14,72,73], we discarded the hypothesis of very weak or no selection on the X as an explanation of the lack of signal observed in the 401 SNP window scan. Additionally, as shown in Garud et al. 2015 and our simulations (**Figure S5**), H12 has power to detect complete hard sweeps. Thus, it is unlikely that H12 has missed such signatures on the X.

Comparing the distribution of the 401 SNP window size in terms of base pairs, we found that the average window length (bp) is ~ 1.5 times larger on the X than on the autosomes (p-val < 2.2e-16, one-sided Wilcoxon rank sum test; **Figure S6**). Moreover, the increased recombination rate on the X exacerbates differences in window sizes in terms of centimorgans, as the size of the footprint of selection (bp) decreases with higher recombination, resulting in a stronger and faster LD decay (**Figure 4, Figure S2)**. Therefore, it is possible that 401 SNP windows are too large to effectively detect selection on the X chromosome.

To be able to define H12 analysis windows that are more comparable in terms of base pair length between the X chromosome and the autosomes, we defined smaller windows for the X chromosomes with an average length ~10kb (**Figure 5**). More concretely, we used the autosomal and X chromosome S/bp median values obtained from the DGRP data to redefine the number of SNPs per window in our H12 scan. The median S/bp in the autosomes of the DGRP data is 0.0345, which means that 401 SNP windows correspond to a window length of *L* = 11,623bp. For a recombination rate of 5e-7cM/bp, sweeps with S ≥ 0.05% are likely to generate a signature that extend approximately *L* = 11623bp (*L* ≈ *s*/[log(*N_e_s*) *ρ*]). The median S/bp on the autosomes is 0.0227, hence, if we took 401 SNP windows as before, the windows in terms of base pairs would be approximately 17,665 bp long. Furthermore, for *ρ*=5e-7cM/bp and *N_eX_* = 0.75*N_eAuto_*, sweeps with *s~* 0.1% or greater would be observed in windows of this length. For higher recombination rates, as in the case of the X chromosome (**Figure S2**, [68]), only selective sweeps with approximately *s>* 0.1% would be observed. Therefore, to make the H12 analysis windows of the autosomes and the X chromosome more comparable, we defined the X chromosome windows by 0.0227 × *L* ≈ 265 SNPs, where *L* is the autosomal window distance calculated previously. Furthermore, for the autosomes, we took 401 SNP windows, which we randomly down sampled to 265 SNPs, making our analysis windows equivalent in terms of numbers of SNPs.

**Figure 5.**
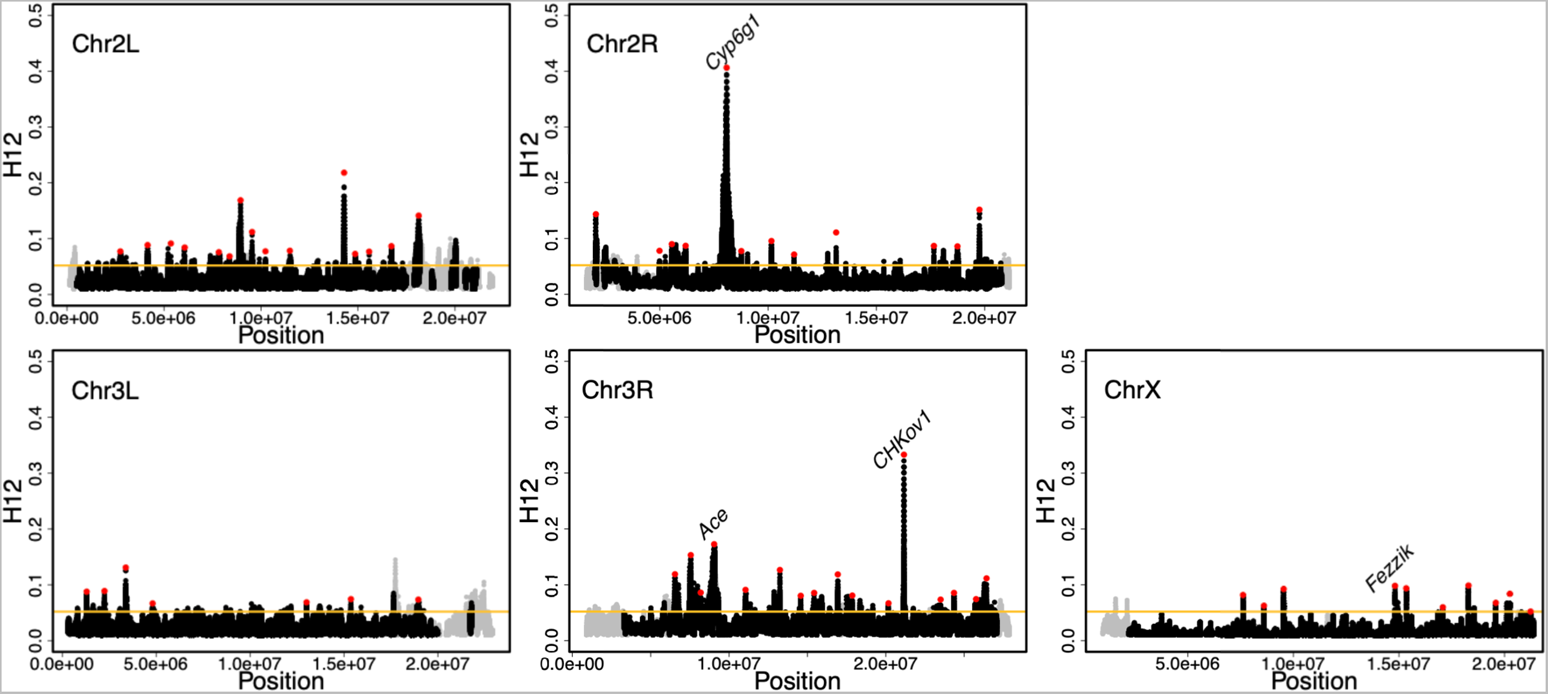
H12 scan in the DGRP data. H12 scan in DGRP data for four autosomal arms and the X chromosome. For the autosomal scan, each data point represents an H12 value in a 401 SNP window down sampled to 265 SNPs. For the X chromosome, windows of 265 SNPs were used. Regions with recombination rates lower than 5×10^−7^ cM/bp were excluded from the scan and are denoted in grey. The red data points denote the top 50 and top 10 autosomal and X chromosome peaks, respectively. The four positive controls (*Ace*, *Cyp6g1*, *CHkov1* and *Fezzik*) are highlighted in the scan. The golden line represents the 1-per-genome FDR line calculated under admixture Model 1 and a recombination rate of 5×10^−7^ cM/bp (**Methods**).

#### 2.3. Softness of sweeps on the X versus autosomes

To gain intuition on the haplotype structure of the top peaks of the autosomes and the X chromosome, we visualized their haplotype frequency spectra (**Figure 6a**). We also visualized the haplotype frequency spectra of hard and soft, partial, and complete sweeps from simulations (**Figure 6b**). Several peaks on the autosomes have multiple haplotypes at high frequencies, consistent with signatures of soft sweeps; whereas more peaks on the X have haplotype frequency spectra that resemble partial and hard sweeps.

**Figure 6.**
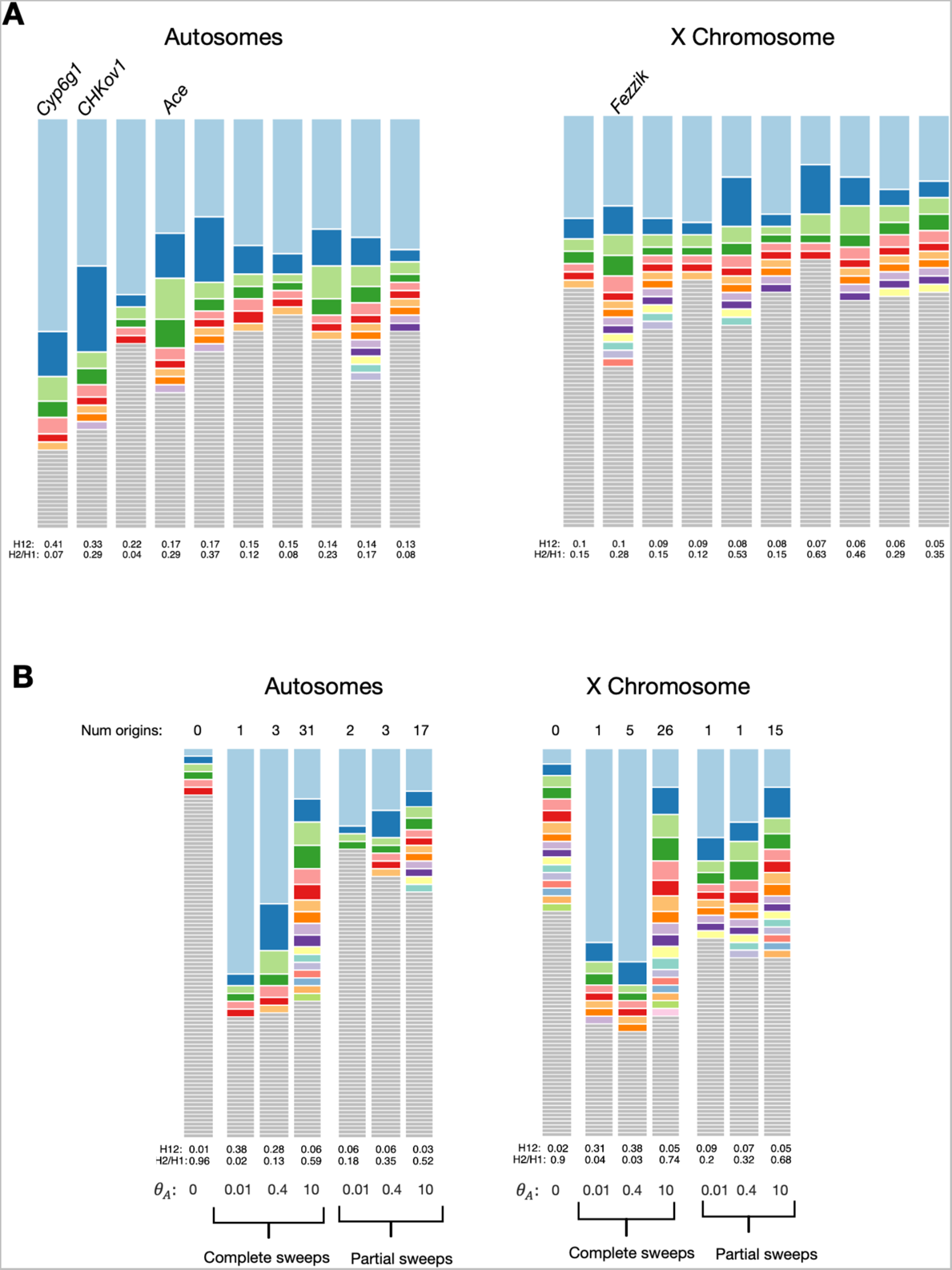
Haplotype frequency spectra for top DGRP peaks and simulated hard and soft sweeps. (**A**) Top ten autosomal and X chromosome peaks. The analysis window with the highest H12 value is plotted and peaks are ordered from highest to lowest H12 (left to right). Each colored bar represents a distinct haplotype, and the size of the bar corresponds to its frequency in the sample. Gray bars indicate singletons. (**B**) Expected haplotype frequency spectra in simulations of a neutral constant *N_e_* model and selective sweeps from de novo recurrent mutations. These simulations were randomly chosen. Compared are three complete sweeps and three partial sweeps with partial frequency PF=0.5. The adaptive mutation rate θ_A_ is varied to be 0.01, 0.4 and 10 with *h*=0.5.

To determine whether the top peaks in our scan were more likely generated by hard or soft sweeps we computed H2/H1 [39], which in conjunction with high H12 values can differentiate hard and soft sweeps [39,50]. H2/H1 is the ratio of haplotype homozygosity excluding the most frequent haplotype (H2) and standard haplotype homozygosity (H1). Given that in a hard sweep a single haplotype is found at a high frequency, hard sweeps are expected to have low H2/H1 values. As sweeps become softer, more haplotypes are present at substantial frequencies, increasing H2/H1 monotonically with the softness of the sweep [50]. Thus, as proposed by Garud et al 2015, H12 together with H2/H1 can distinguish whether a sweep is more likely to be hard or soft.

We used an approximate Bayesian computation (ABC) approach to differentiate the likelihood that a given (H12, H2/H1) pair is generated by a hard or a soft sweep model (**Figure 7**). This likelihood is given by Bayes factors defined as BF= P(H12_obs_, H2_obs_/H1_obs_ | soft sweep)/ P(H12_obs_, H2_obs_/H1_obs_ | hard sweep), where H12_obs_ and H2_obs_/H1_obs_ were computed from the DGRP data. Hard sweeps have BF ≤ 1 while soft sweeps have BF > 1, with stronger evidence for soft sweeps given for BF >> 1.

**Figure 7.**
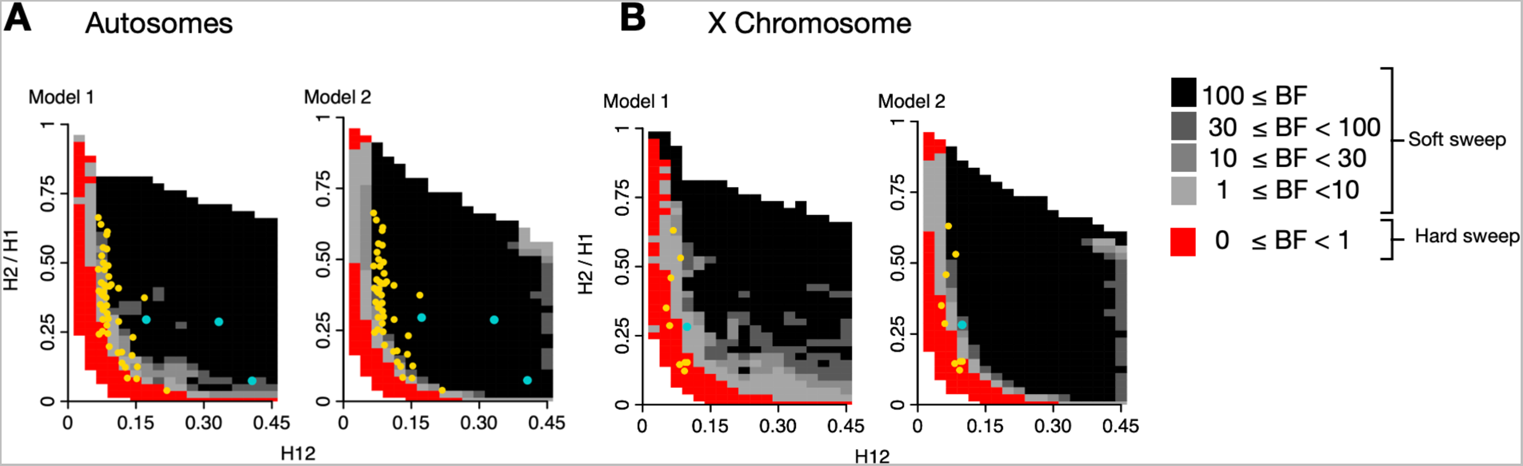
Expected H12 and H2/H1 parameter region for hard and soft sweeps for the autosomes and X chromosome under two variations of the Duchen et al. admixture demographic models. In Model 1, North American *N_e_*=1,110,000, and in Model 2, *N_e_*=15,984,500. We obtained the expected H12, H2/H1 parameter region expected for hard and soft sweeps calculating BFs for a grid of H12 and H2/H1 values. We calculated BFs by computing the ratio of soft and hard sweeps obtained from simulations found within a Euclidean distance of 0.1 of an (H12, H2/H1) pair. The (H12, H2/H1) parameter region that is more likely to be generated by hard sweeps is indicated in red (BF<1). The parameter region that is more likely to represent soft sweeps is shown in gray (BF ≥ 1), where the darker the gray the higher the likelihood of a soft sweep (BF ≥ 30). The yellow points highlighted correspond to the top 50 autosomal peaks (A) and top 10 X chromosomal peaks (B). The blue dots are the positive controls at *Ace*, *CHKov1*, *Cyp6g1* and *Fezzik*. We used 401 SNP windows down sampled to 265 for the autosome simulations and 265 SNP windows for the X chromosome simulations

We simulated hard and soft sweeps under the admixture Models 1 and 2, drawing the selection coefficient (*s)* and age of the sweep *T_E_* from uniform prior distributions *s* ~ *U*[0,1] and *T*_2_ ~ *U*[0, 1034] × 4*N_e_* (Methods). Hard sweeps were generated under *θ_A_* = 0.01 and soft sweeps under *θ*_!_ = 10 [19,20]. We ran 5×10^5^ hard and soft sweep simulations for each admixture model using the coalescent simulator *msms* [74] for a constant recombination rate of *ρ* = 5 × 10^35^cM/bp. Given that the average recombination rate is higher on the X than on the autosomes (**Figure S2,** [68,71]), we also simulated hard and soft sweeps on the X with *ρ* = 1 × 10^35^ cM/bp and *ρ* = 1 × 10^36^ cM/bp (**Figure S7**). With *msms* we are able to run the high number of simulations required for ABC while accounting for demography, which is not computationally feasible using SLiM (see Discussion). Additionally, we performed ABC for a constant *N_e_*=2.7×10^6^ model running 5×10^4^ hard and soft sweep simulations on SLiM, incorporating male hemizygosity and including partial frequency after selection ceased (*PF*~*U*[0,1]) and dominance (*h*~ *U*[0,1]) in our nuisance parameters (Methods) (**Figure S8**).

The panels in **Figures 7, S7 and S8** show the BFs calculated from our simulations for a grid of H12 and H2/H1 values. In both Model 1 and Model 2, we observe a significantly higher proportion of Hard/Soft Sweeps on the X chromosome than on the autosomes (one-tailed Exact Fisher’s test p-val=0.001 and p-val = 2.75×10^−5^ for Model 1 and Model 2, respectively). For X chromosome simulations with higher recombination rates, we still observe a significantly higher proportion of Hard/Soft sweeps compared to autosomes, although the number of sweeps classified as hard decreases when recombination is high (**Figure S7**). Additionally, for the constant *N_e_* model, in which *PF* and *h* were also treated as nuisance parameters, we also observe an enrichment of hard sweeps on the X (one-tailed Exact Fisher’s test p-val=0.0001). This suggests an enrichment of hard sweeps on the X chromosome that is robust to demography.

## Discussion

It has been suggested that due to male hemizygosity, the X chromosome experiences more efficient selection as well as an accelerated rate of evolution compared to autosomes [13,56,57,65,75]. However, despite widespread interest in the evolutionary forces shaping the X chromosome and the autosomes, our understanding of the tempo and mode of adaptation in natural populations is still forming with the emergence of new datasets [51] and statistical methods to detect selection [39,76–81]. In this study, we found that sweeps are, on average, expected to be harder on the X than on autosomes in a variety of simulated evolutionary scenarios. Confirming this prediction, we found evidence supporting an enrichment of hard sweeps on the X chromosome of North American *D. melanogaster* in the DGRP dataset.

Our finding that hard sweeps are enriched on the X in *D. melanogaster* is in accordance with recent work. Recently, Muralidhar and Veller [21] showed that dominance shifts can lead to softer sweeps on the autosomes. They suggested that because the X is hemizygous in males and thus cannot experience dominance shifts in males, soft sweeps from SGV may be less common on the X, which we confirm. Additionally, recent work in great apes showed that deep dips in diversity on the X chromosome are more consistent with hard sweeps as compared to soft sweeps [75], providing empirical support for the notion that hard sweeps may in fact be common on the X. Finally, it has also been suggested that *D. mauritiana’s* X chromosome experiences more hard than soft sweeps [82].

Consistent with Singh et al. [65], which found that neutral demography alone cannot account for dips in diversity on the X relative to the ancestral African population and autosomes, we found evidence for selective sweeps on the X. To do so, we extended the autosomal H12 scan we performed in previous work [39,52], in which we found evidence for abundant soft sweeps.

Due to the significant reduction of diversity on the X chromosome (**Figure 4**), we defined H12 windows for the X and autosomes such that they are comparable both in terms of length and SNP density. We note that a window defined with a fixed number of base pairs instead of SNPs can potentially result in noisier scans because of random dips in diversity due to drift and background selection. By contrast, defining windows with a fixed number of SNPs ensures that H12 does not co-vary with the number of SNPs available per window. Down sampling the number of SNPs for the autosomal windows from 401 to 265 did not alter the results obtained with the original 401 SNP windows (**Figure 5 and Figure S3**), including detection of known soft sweeps at *Ace, Cyp6g1,* and *CHKov1.* Moreover, using 265 SNP windows on the X allowed us to recover the signal near *Fezzik*, which was not possible with 401 SNP windows (**Figure S3**). In the future, our window approach may be useful for studies comparing populations with substantial differences in their diversity levels.

The recovery of the *Fezzik* locus in our scan was confirmatory given that previous work has shown that the *Fezzik* enhancer has experienced directional positive selection in derived populations of *D. melanogaster* compared to populations from sub-Saharan Africa [14,49,83]. The *Fezzik* gene has been shown to affect tolerance to cold and insecticides [49] and is also thought to be involved in ecdysteroid metabolism [84] and oxidoreductase activity [85]. However, whether this locus experienced a hard or soft sweep was previously unknown. Interestingly, this peak was classified as soft in our ABC analysis. This result may be consistent with the fact that Glaser-Schmitt et al. [14] showed that a SNP located within the *Fezzik* enhancer is likely under balancing selection as a result of sexually antagonistic forces and temporally fluctuating selection acting on *Fezzik* expression in males and females [14]. Their results predict that the variant under selection is likely female beneficial and dominant, although varying dominance might be involved. In our simulations of sexual antagonism (**Figure 3**), we observed that when mutations are beneficial in females and deleterious in males, the likelihood of soft sweeps increases with dominance for both the X and the autosomes.

The *Fezzik* case example provides some biological insight into the underlying mechanisms that could generate soft sweeps on the X. However, being able to accurately distinguish which scenario is driving adaptation is challenging, as the observed signatures could be a result of multiple evolutionary processes. For example, other scenarios that could explain soft sweeps on the X include adaptation occurring by recurrent recessive beneficial mutations. Similarly, as we show in our simulations, hard sweeps on the X could be the result of partially dominant mutations or adaptation to a change in the environment through constant dominance or dominance shifts (**Figures 1–3**). Additional simulations that incorporate more variations to the model of sexual antagonism such as varying dominance, differences in the magnitude of selection between males and females and temporally fluctuating sex-dependent selection, and variations on demography are needed to better understand the effect of more complicated evolutionary scenarios on the signatures of selection.

We acknowledge that the demographic models used for our ABC analysis are only estimates of the North American *D. melanogaster’s* population history and may not fully capture the complexity of this admixed population. To the best of our knowledge, a neutral model that fits the data in terms of S/bp, Pi/bp and long-range LD is not currently available. We therefore used two admixture models proposed in [52] since these were shown to provide a better fit to the data than previous models [39,70,86,87]. Our results showed evidence for an enrichment of hard sweeps on the X chromosome of the DGRP data regardless of the underlying demographic model tested (**Figure 7**) or increased recombination rate of the X (**Figure S2**). In addition, we tested a constant *N_e_* model with male hemizygosity incorporated and found similar results (**Figure S8**). Future work that explores the multidimensional parameter space of *D. melanogaster’s* demographic history in search of a model that fits multiple genome-wide statistics to the data would greatly benefit the field. We note that it is possible that to obtain a model that provides a good fit across population genetic statistics, more complex scenarios, such as the effect of seasonal fluctuations [88,89] and selection might need to be included.

Ideally, our simulations of X chromosome and autosome evolution would simultaneously incorporate both male hemizygosity and admixture. However, doing so is currently a challenge since forward in time simulators such as SLiM [53], that provide the flexibility to model sex chromosome evolution, dominance shifts, and sexual antagonism, cannot simultaneously handle selection and the large effective population sizes and complex demography of *D. melanogaster* populations [53]. On the other hand, coalescent simulators like *msms* [74] can model complex admixture events for large *N_e_* but cannot model male hemizygosity. Approaches such as rescaling and tree sequence recording have been shown to increase the computational efficiency of forward in time simulators [54,90,91]. Nonetheless, rescaling has not been proven to maintain the genetic diversity of the original model when complex demography and selection are simulated together. Simulating the admixture models in SLiM with the parameters in their original scale would be computationally unfeasible. To address this issue, we modeled the X chromosome by reducing its effective population size to ¾ of the *N_eAuto_* and ran our admixture models using *msms* [74]. Additionally, we ran a constant *N_e_* model in SLiM, incorporating male hemizygosity.

Although we now have evidence for a handful of species in which hard sweeps are more common on the X than the autosomes, it remains to be seen if this is generically true of all species. If hard sweeps are more prevalent on the X than on the autosomes across populations, future work could seek to answer whether hard sweeps are important in driving sexual dimorphism and speciation, where the X chromosome has been shown to play a significant role [1,2,4,15]. Moreover, continuing to study the signatures of selection on the X and the autosomes will further increase our understanding on how demographic forces, as well as other evolutionary variables, such as dominance, differentially affect the X and the autosomes.

## Methods

### Data Processing

We used the publicly available Drosophila Genome Nexus (DGN) dataset (Lack et al. 2015), which includes 205 DGRP strains from Raleigh (RAL), North Carolina and 197 DPGP3 strains from Zambia (ZI). These data can be downloaded at www.johnpool.net.

In order to avoid false positives resulting from IBD from closely related strains, we removed strains with genome-wide IBD levels greater that 20% with at least one other strain. These correspond to 8 ZI strains and 27 RAL strains: ZI397N, ZI530, ZI269, ZI240, ZI218, ZI207, ZI523, ZI86,RAL-385, RAL-358, RAL-712, RAL-399, RAL-879, RAL-355, RAL-810, RAL-350, RAL-832, RAL-882, RAL-306, RAL-799, RAL-801, RAL-859, RAL-907, RAL-790, RAL-748, RAL-336, RAL-850, RAL-365, RAL-786, RAL-730, RAL-861, RAL-59, RAL-646, RAL-812, and RAL-787. This resulted in a total of 178 RAL strains and 189 ZI strains.

The North Carolina DGRP dataset consists of data from flies that were extensively inbred to obtain mostly homozygous genomes. Nevertheless, this inbreeding process left tracts of residual heterozygosity, which, in some cases, are substantial. These tracts of residual heterozygosity were treated as missing data, and, if not accounted for, can give false H12 signals. To reduce the inflation of the H12 statistic caused by the remaining IBD and from the masking of heterozygous sites, we down sampled to the top 100 strains with least amount of missing data for each chromosome, separately. Moreover, we required each site to be called in at least 50% of the genome (**Figure S9**).

### Computation of summary statistics

To calculate LD, we used the *R*^2^ statistic in sliding windows of 10kb, iterating by 50bp. We only considered SNPs with alleles frequencies between 0.05 and 0.95. SNPs with missing data were excluded and at least 4 individuals at both SNPSs were required to calculate LD. We then smoothed the LD plots as in Garud et al. 2015 by averaging LD values binned in 20 bp windows until 300 base pairs were reached, after which LD values were averaged in windows of 150 base pairs.

We computed S/bp and Pi/bp in non-overlapping 10kb windows for the DGRP data. We estimated the mean levels of *θ*_8_ and the corresponding confidence intervals by bootstrapping. We performed 1000 bootstrap replicates per parameter estimated and constructed the 95% confidence intervals corresponding to each bootstrapped distribution.

### SLiM Simulations

We used SLiM 3.7 [53] to simulate autosomal and X chromosome evolution. For simplicity we simulated a constant *N_e_*=10^6^ demographic model under 3 different scenarios: recurrent beneficial mutations, dominance shifts and sexual antagonism. Since SLiM is a forward-in time simulator, simulating large population sizes is computationally intensive. We found that for population sizes greater than 5×10^5^ simulations become intractable. To make our simulations feasible, we performed rescaling on our model parameters. To do so, we followed Algorithm 1 from Uricchio and Hernandez [90] with a rescaling constant of Q=50 [90,91]. Both algorithms proposed by these authors closely maintain the levels of genetic variation of the non-rescaled population in a constant *N_e_* model of *D. melanogaster,* as long as selection is not too strong (*s<0.1)*.

We kept track of the number of distinct mutational origins in our simulations and defined a sweep as soft if a sample had two or more distinct mutational origins. For the dominance shift simulations, we kept track of the number of mutations present at the time of the shift. If there were no mutations present, we labeled the simulation as “no standing variation”. If there were mutations at the time of the shift but then the mutations were lost, we called it “lost sweep”. We simulated sexual antagonism in two ways: male advantage with female disadvantage and male disadvantage with female disadvantage. The deleterious mutation in each of these scenarios was defined as *s_d_* = −*ks_b_*, with *k* a scalar that modulates the strength of selection.

### H12 scan

We ran a genome-wide scan using sliding windows of 401 SNPs down sampled to 256 SNPs for the autosomes and 256 SNPs for the X chromosome, iterating by intervals of one SNP between window centers. To avoid false peaks driven by high missingness, we excluded windows with more than 10% of missing data from haplotype clusters and considered them as singletons in the scan.

We called peaks by identifying the windows with H12 values above the H12_o_ critical value. We defined H12_o_ to be the false discovery rate (FDR), which corresponds to the 10^th^ highest H12 value obtained from 10 times the number of independent analysis windows in the data (~100,000) neutral simulations. We obtained H12_o_ for each admixture model as well as a constant *N_e_*=2.7×10^6^ model and chose the highest H12_o_ as our critical value (H12_o_ =0.052).

We grouped the consecutive windows above the threshold into a single peak and obtained the highest H12 value to represent the value of the peak. We then iterated through the identified peaks, from highest to lowest H12 value, and excluded the peaks found within 500 kb of the center of the peak under inspection. This avoided the identification of peaks belonging to the same selective event. We finally masked peaks found in regions of low recombination (< 5×10^−7^ cM/bp) identified using the Comeron et al. (2012) crossover map.

### Approximate Bayesian Computation (ABC) to classify hard and soft sweeps

We used ABC to compute the likelihood that a soft or a hard sweep model generates a pair of (H12,H2/H1) values. We simulated hard sweeps with *θ_A_* = 0.01 and soft sweeps with *θ*_!_ = 10. To be able to run a large number of simulations, we used the coalescent simulator *msms* [74]. We ran a total of 5×10^5^ simulations for both the hard and soft sweep models for the two admixture models proposed by Garud et al. (2021). These models are variations of the Duchen et al 2013 admixture model and were fitted to the autosomal DGRP data in terms of the summary statistics S/bp, Pi/bp and H12 while accounting for admixture events in North American *D. melanogaster.* Because *msms* does not have the option to simulate sex chromosome evolution, we simulated the X chromosome by downscaling the effective population sizes by ¾.

We drew the values of the nuisance parameters selection strength (*s)* and age of sweep (*T_E_*) from the following prior distributions: *s* ~ *U*[0,1] and *T*_2_ ~ *U*[0, 1034] × 4*N_e_*. We then calculated Bayes factors for a grid of (H12, H2/H1) values by taking the ratio of the number of soft sweep and hard sweep simulations with an Euclidean distance <0.1 from each (H12, H2/H1) data point from our H12-H2/H1 grid.

Additionally, to incorporate male hemizygosity into our X chromosome simulations, we modeled a constant *N_e_*=2.7×10^6^ population on SLiM. As before, hard sweeps were simulated under *θ*_!_ = 0.01 and soft sweeps under *θ*_!_ = 10. Due to the large *N_e_* of the model, we downscaled our simulation parameters by a factor of Q=50 [90] and used tree sequence recording to add neutral mutations to our model and perform recapitation with *msprime* [92] and *pyslim* [54], making the simulations computationally tractable. In addition to the hyperparameters *s* and *T*_2_, these simulations include partial frequency after selection ceased (*PF*~ *U*[0,1]) and dominance (*h*~ *U*[0,1]).

## Code Availability

Code used to process and analyze the data is available at: https://github.com/garudlab/SelectiveSweeps_Xchr_vs_Auto

## Acknowledgements

We sincerely thank Dmitri Petrov for his insights, which inspired this work. We also sincerely thank Kirk Lohmueller for his feedback on this work at several stages and Alison Feder for her feedback on the manuscript. We thank the Garud and Lohmueller labs for their feedback on the manuscript. MH was supported by the Systems and Integrative Biology Training Grant (NIH-NIGMS 5T32GM008185-33) and the Training Grant in Genomic Analysis and Interpretation (NIH T32HG002536).

## Notes

### Competing Interest Statement

The authors have declared no competing interest.

