## Supplementary Figures and Table S1 for "Enrichment of hard sweeps on the X chromosome in *Drosophila melanogaster*"

### Supplementary Material

**A) Recurrent Adaptive Mutations**

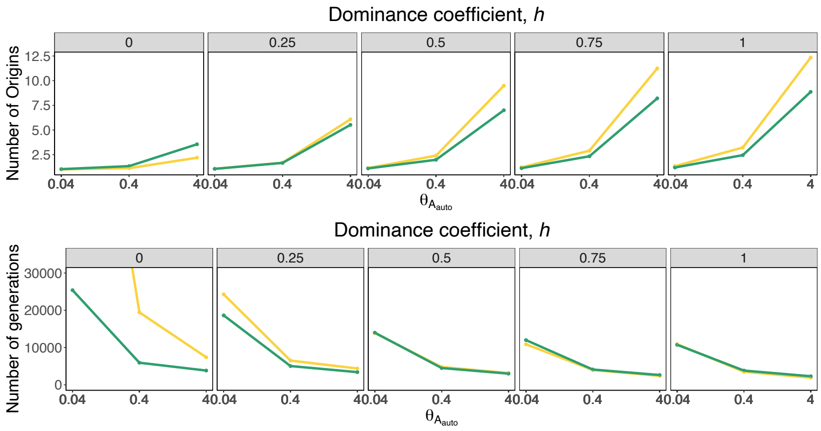


**B) Sexual Antagonism–Female Disadvantage**

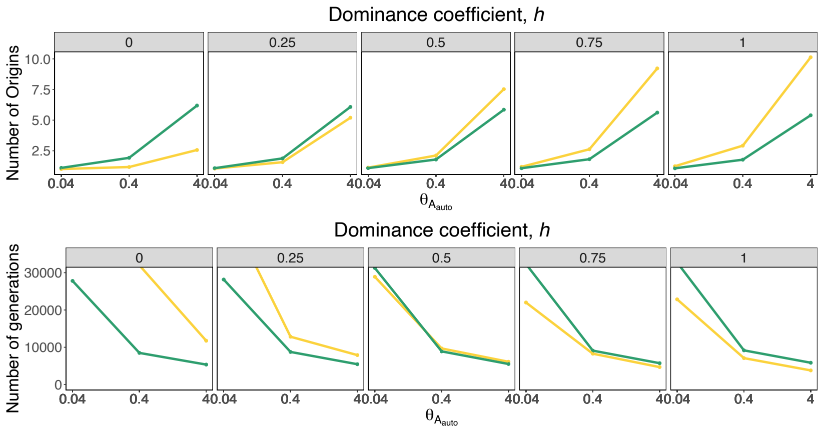


**C) Sexual Antagonism–Male Disadvantage**

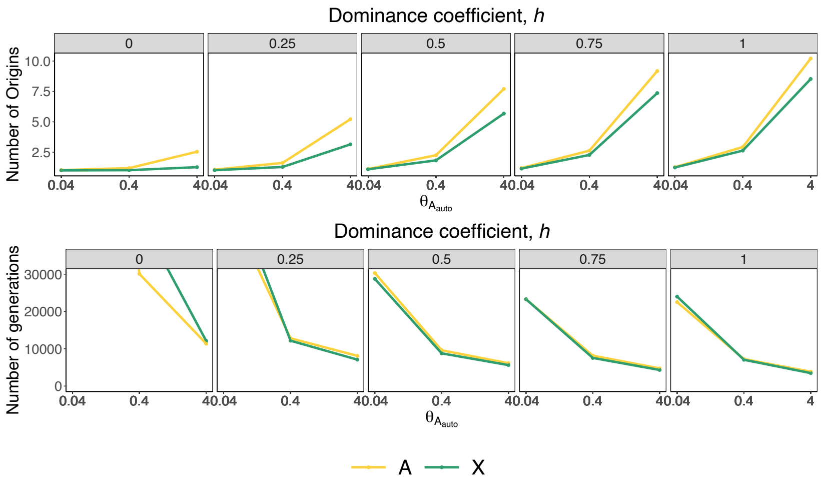

**Figure S1. Average number of origins and average number of generations until fixation in three simulated scenarios.** The average number of origins and generations until fixation were computed across 1,000 simulations of each evolutionary scenario per combination of parameters. The yellow lines correspond to the autosomal values and the green lines to the X chromosome. The scenarios depicted in the figure include (**A**) Adaptation through recurrent *de novo* mutations, (**B**) Sexual antagonism with female disadvantage and (**C**) Sexual antagonism with male disadvantage.

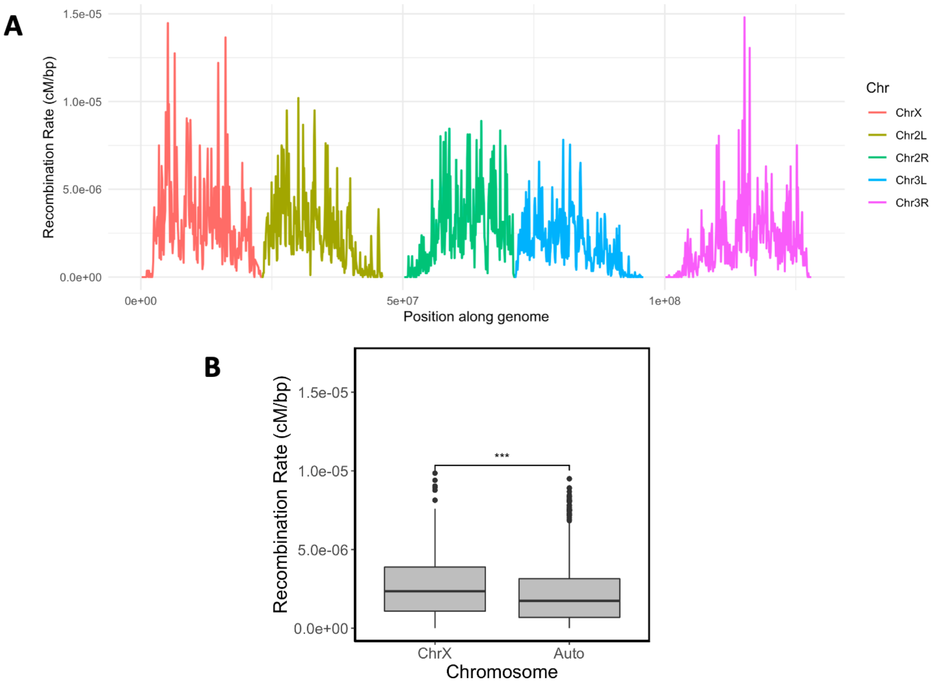


**Figure S2. Recombination rates along the genome of D. melanogaster, using the Comeron et al. (2012) crossover map.** (**A**) Recombination rates along the genome colored by chromosome. (**B**) Distribution of X chromosome and autosomal recombination rates. The distributions of recombination rates on the X chromosome is significantly higher than on that of the autosomes (one-sided Wilcoxon rank sum test p-val =8.685e-06).

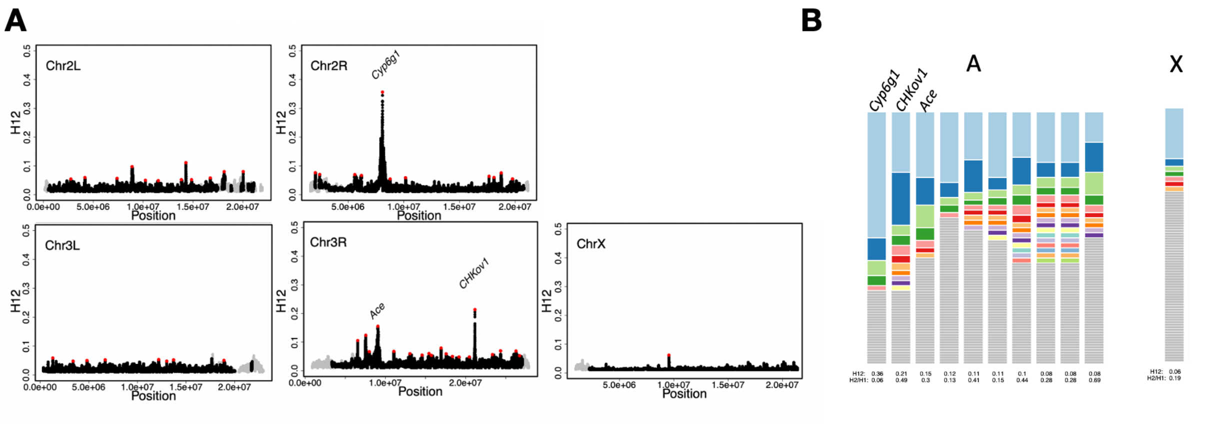


**Figure S3**. **H12 scan with 401 SNP windows for four autosomal arms and the X chromosome.** (**A**) H12 scan in DGRP data for four autosomal arms and the X chromosome using 401 SNP windows. Gray points correspond to genomic regions with recombination rates lower than 5x10-7 cM/bp. The red data points denote the top 50 autosomal peaks and the only X chromosome peak detected in the 401 SNP window scan. (**B**) Haplotype frequency spectra for the top 10 autosomal peaks and the X chromosome peak. Each colored bar represents a distinct haplotype, and the size of the bar corresponds to its frequency in the sample. Gray bars indicate singletons. The three autosomal positive controls, Ace, Cyp6g1, CHkov1 are highlighted in the figure.

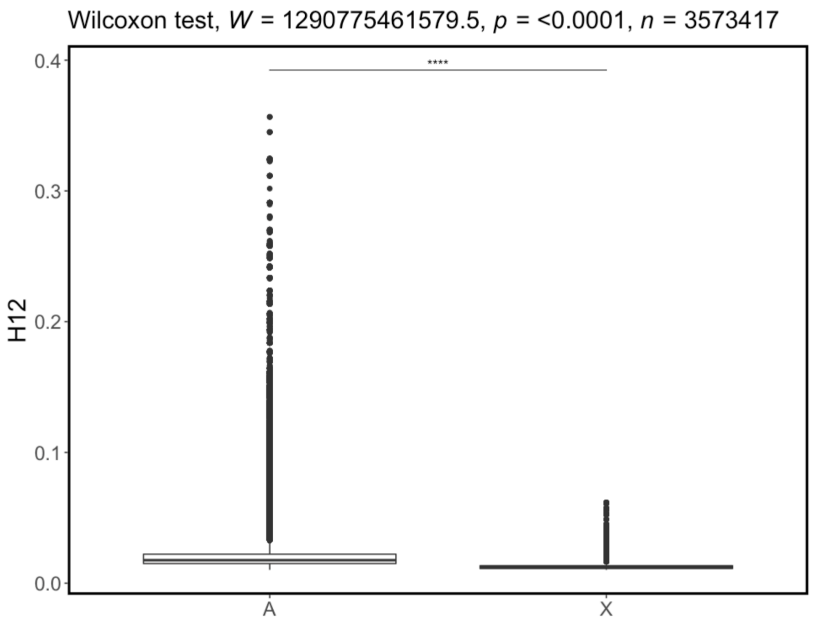


**Figure S4. Distribution of H12 values for 401 SNP windows in the autosomes and X chromosome of *D. melanogaster*.** A Wilcoxon rank sum test on the distribution of the autosomal and X chromosome H12 values shows that the distributions are significantly different with p-val<2.2e-16.

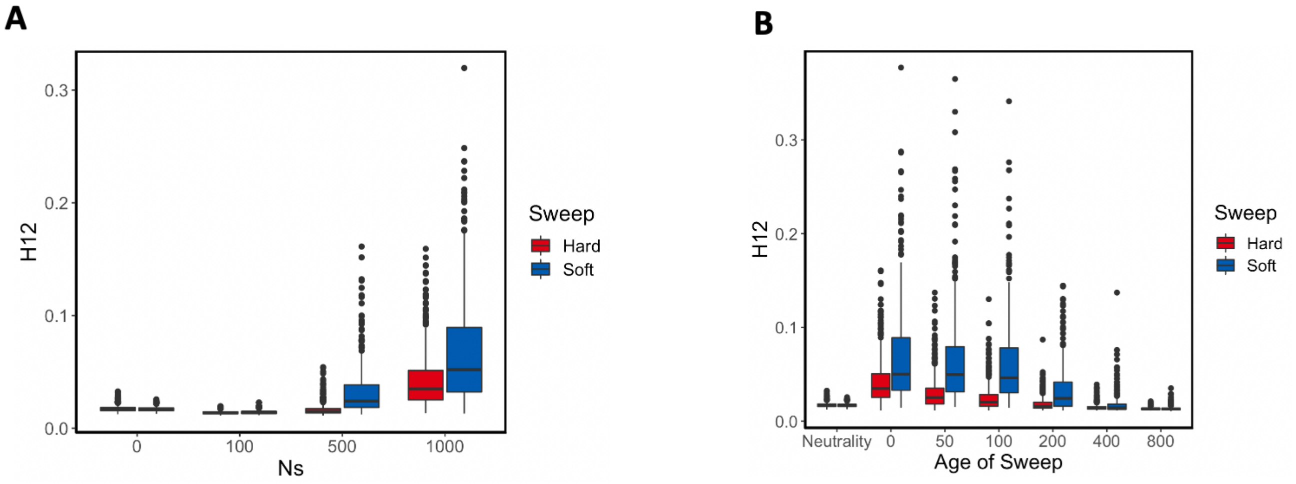


**Figure S5. Ability of H12 to detect complete hard and soft selective sweeps**. Simulations of complete hard and soft sweeps for varying selection strengths ($N_e s$) and increasing age of the sweep. The age of the sweep is given in number of generations after fixation, when the selection strength ceased. (**A**) Distribution of H12 values for hard (red) and soft (blue) selective sweeps for varying strengths of selection s at the time of fixation. H12 can distinguish selection from neutrality when selection is sufficiently strong. (**B**) Distribution of H12 values for hard and soft sweeps with s=0.1, for increasing number of generations after fixation. H12 can detect selection for sweeps that are not too old (age <200 generations).

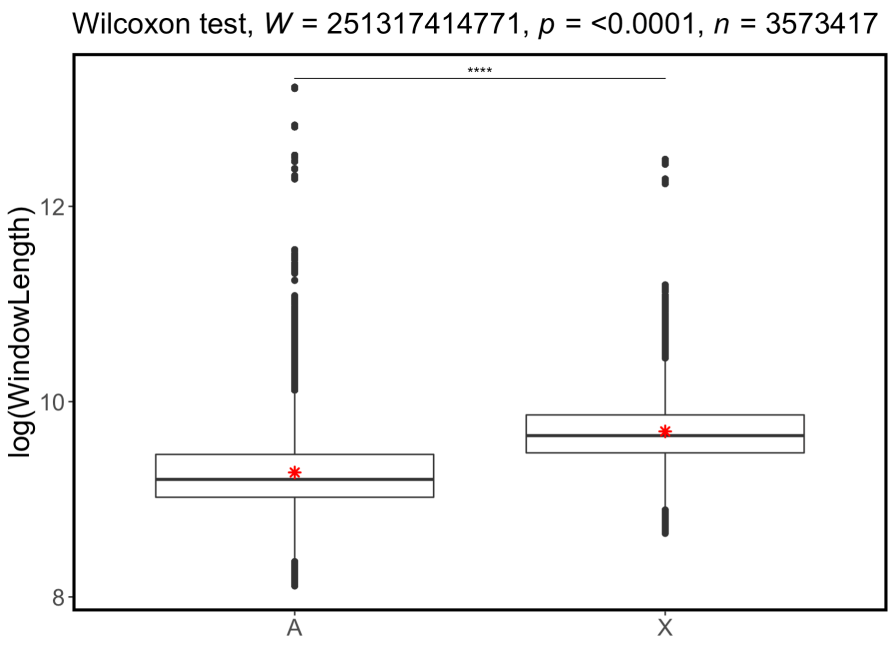


**Figure S6. Window length (bp) distributions in H12 scan performed with 401 SNP windows**. Distribution of the log of the window lengths for the autosomes (A) and the X chromosome (X). The red star indicates the mean window length in the autosomes and the X, respectively. Autosomes have significantly lower window lengths than the X chromosome in the 401 SNP window scan (p-val<2.2e-16, one-sided Wilcoxon rank sum test).

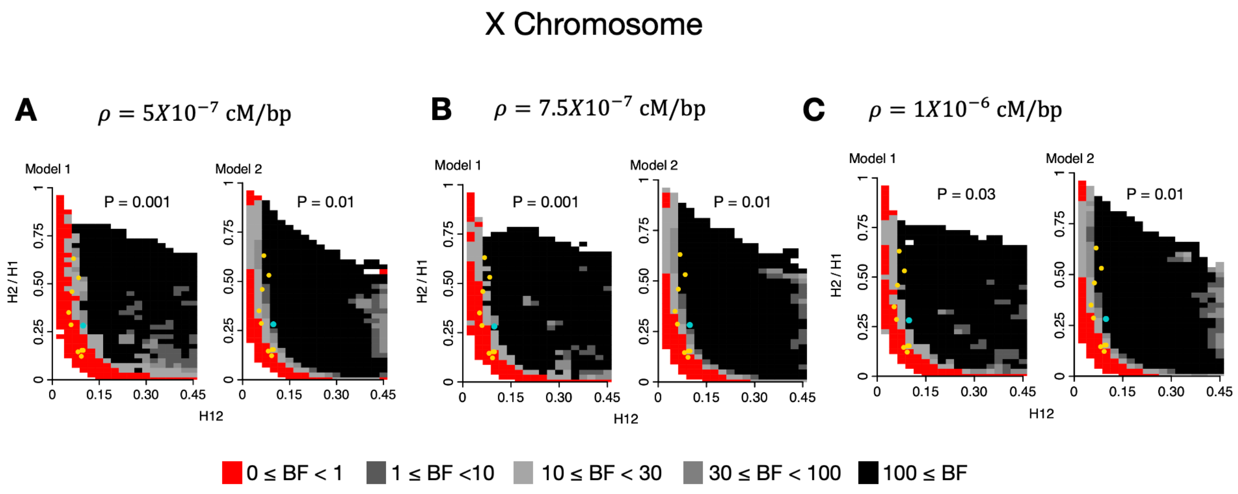


**Fig S7. Expected H12 and H2/H1 parameter region for hard and soft sweeps for the X chromosome for varying recombination rates.** A total of $10^5$ hard and soft sweep simulations were performed for the following recombination rates: $\rho = 5x10^{-7}$ cM/bp (**A**), $7.5x10^{-7}$ cM/bp (**B**) and $1x10^{-6}$ cM/bp (**C**). Regions shown in red provide support for hard sweeps whereas regions in gray show support for soft sweeps. Each panel shows a p-value corresponding a one-sided exact fisher test comparing the hard/soft sweep between the X chromosome (scenarios **A-C**) and the autosomes. The number of hard and soft sweeps on the autosomes were obtained from an ABC analysis using $\rho = 5x10^{-7}$ cM/bp (**Figure 7)**. These simulations were done on *msms,* accounting for admixture and setting $N_{eX}=3/4N_{eAuto}$.

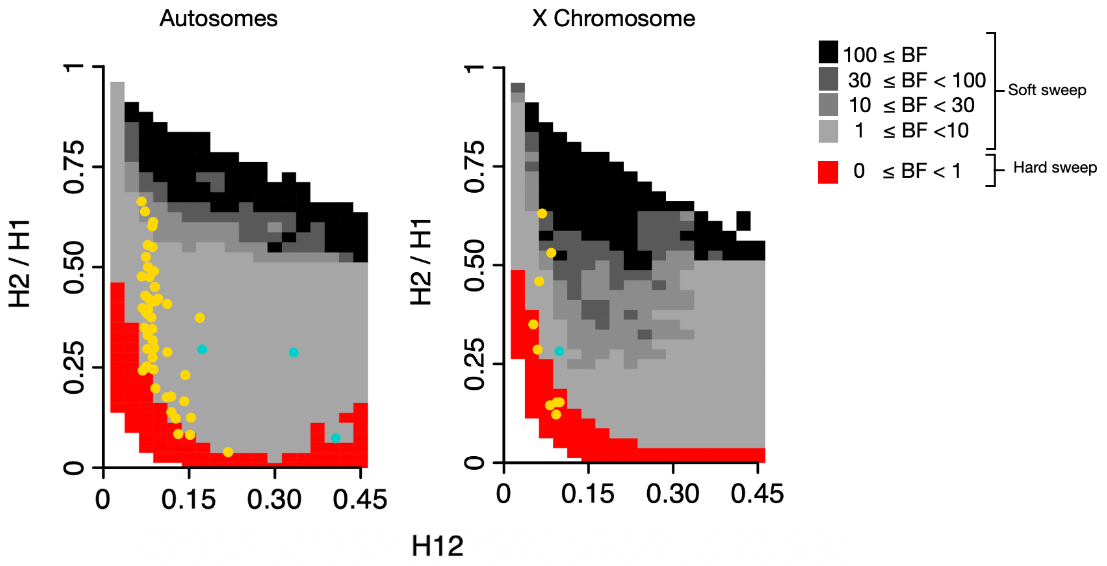


**Fig S8. Expected H12 and H2/H1 parameter region for hard and soft sweeps for the X chromosome and the autosomes in a Constant $N_e$=2.7x10$^6$ model.** ABC analysis to classify H12 peaks as hard and soft. A total of 5x10$^4$ hard and soft sweep simulations were performed using SLiM for each the autosomes and the X chromosome. The regions colored in red show parameter values that are more likely to be generated by soft sweeps, while gray regions correspond to the parameter space that is more likely to be generated by soft sweeps. The points highlighted in the two panels correspond to the top 10 H12 peaks on the X chromosome of the DGRP data with the blue point corresponding to the positive controls. Nuisance parameters were drawn from uniform prior distributions as follows: $s \sim U[0,1]$ *and* $T_E \sim U[0, 10^{-3}] \times 4N_e$, $PF \sim U[0,1]$ and $h \sim U[0,1]$. These simulations were performed on SLiM, modelling the hemizygosity of the X chromosome.

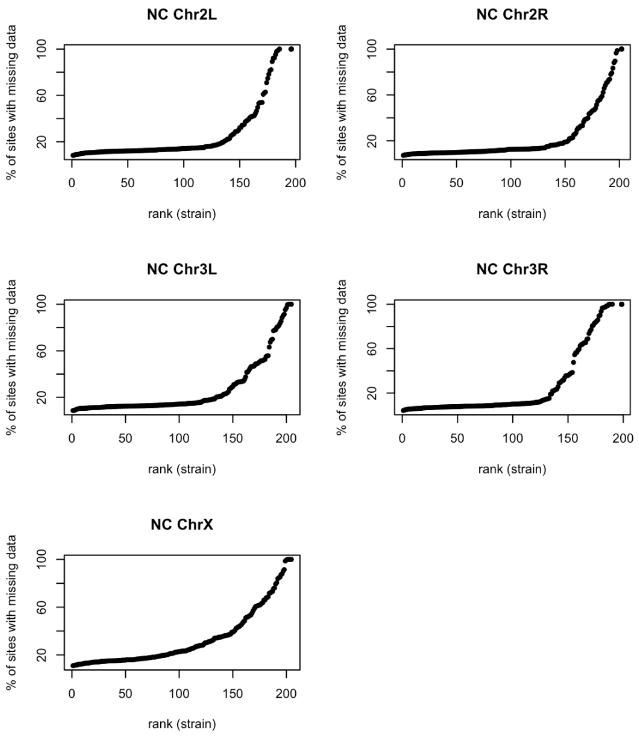


**Figure S9. Number of sites with missing data in each strain in DGRP data**

**Table S1. Demographic parameters for Admixture models from Garud et al. 2021**

Shown are the parameter estimates for the admixture models inferred in [52,69]. The parameters used for both models are explained below.

| | |
|---|---|
| $N_{Aa}$ | Ancestral African population size |
| $sev_A$ | Severity of African bottleneck ($log_{10}$ scale duration/population size) |
| $N_{Ac}$ | Contemporary African population size |
| $N_E$ | European population size |
| $N_{Am}$ | North American population size |
| $T_A$ | Time of bottleneck in Africa ($log_{10}$ scale) |
| $T_{AE}$ | Time of split between African and European populations ($log_{10}$ scale) |
| $T_{adm}$ | Time of admixture between Africa and Europe ($log_{10}$ scale) |
| $prop_{adm}$ | Proportion of European admixture |

**Model 1:**

| $N_{Aa}$ | $sev_A$ | $N_{Ac}$ | $N_E$ | $N_{Am}$ | $T_A$ | $T_{AE}$ | $T_{adm}$ | $prop_{adm}$ |
|---|---|---|---|---|---|---|---|---|
| 5,224,100 | 0.21 | 4,975,360 | 700,000 | 1.11e6 | 5.38 | 5.29 | 3.16 | 0.85 |

**Model 2:**

| $N_{Aa}$ | $sev_A$ | $N_{Ac}$ | $N_E$ | $N_{Am}$ | $T_A$ | $T_{AE}$ | $T_{adm}$ | $prop_{adm}$ |
|---|---|---|---|---|---|---|---|---|
| 5,224,100 | 0.21 | 4,975,360 | 700,000 | 15,984,500 | 5.38 | 5.29 | 3.16 | 0.85 |
